# Migration of Kupffer’s vesicle derived cells is essential for tail morphogenesis in zebrafish embryos

**DOI:** 10.1101/2024.07.04.602018

**Authors:** Jelmer Hoeksma, Jeroen den Hertog

## Abstract

A phenotypic screen of fungal filtrates on developing zebrafish embryos identified metabolites from the fungus *Ceratocystis populicola* to induce ectopic tail formation, including a split notochord and a duplicated caudal fin. Chemical analyses led to the identification of monoterpene alcohols, in particular geraniol, as active compounds inducing ectopic tail formation during a specific 4 h time window during tail bud stage. Embryos from Tüpfel long fin zebrafish (TL) were more susceptible to ectopic tail formation by geraniol than embryos from Wild Indian Karyotpe (WIK) zebrafish, indicating zebrafish strain specificity. RNA sequencing on tail buds of 15-somite stage embryos revealed downregulation of essential genes of the retinoic acid signaling pathway and differential expression of *cyp26a1* and *fgf8a* and downstream *hox*-genes was validated. Time-lapse imaging revealed that Kupffer’s vesicle derived cells failed to migrate shorty after Kupffer’s vesicle collapse upon geraniol treatment and these cells failed to merge with progenitors from the tail bud. Instead, these cells contributed to an ectopic tail, expressing markers for presomitic mesoderm, somite and notochord tissue. Taken together, our data suggests that Kupffer’s vesicle cells harbor tail progenitor capacity, and proper migration of these cells is essential for normal tail morphogenesis.

**Summary Statement:** Inhibition of Kupffer’s vesicle derived cell migration affected tail morphogenesis and resulted in ectopic tail formation in zebrafish embryos.

## Introduction

Zebrafish (*Danio rerio*) is a versatile and powerful vertebrate model to investigate development and disease using genetic screens and high throughput small molecule screens. Previously, we reported a phenotypic screen of 10,207 fungal filtrates using developing zebrafish embryos. This screen led to the identification of biologically active fungal compounds, including compounds which are routinely being used in the clinic (Hoeksma et al., 2019). In addition, we found that 25.2% of the active fungal filtrates induced tail defects. In a follow-up study, we focused on this category and identified the fungal compound cercosporamide as a BMP-signaling inhibitor, based on the dorsalized phenotype it induces in zebrafish tails (Hoeksma et al., 2020), demonstrating that fungal filtrates interfering with tail morphogenesis harbor compounds with interesting biological activities.

Many signaling pathways are activated during zebrafish tail morphogenesis, which together orchestrate proliferation, cell transitions and cell movements required to establish a fully functional structure. Crucial are the formation and tight regulation of morphogen gradients in all three dimensions. The bone morphogenetic protein (BMP) gradient is essential for dorsoventral patterning of the zebrafish tail (Row and Kimelman, 2009). BMP signaling inhibitors such as dorsomorphin and DMH-1 result in a dorsalized phenotype including loss of ventral fins, as does cercosporamide that we identified previously (Hao et al., 2010; Hoeksma et al., 2020; Yang and Thorpe, 2011; Yu et al., 2008). Similar dorsoventral patterning defects are observed in loss-of-function mutants of key genes of the BMP-signaling pathway, including activin receptor-like kinase 2 (alk2) (in zebrafish also known as alk8 or lost-a-fin) (Bauer et al., 2001), its ligand, bmp2 (Kishimoto et al., 1997) or intracellular factors such as smad5 (Kramer et al., 2002). In contrast, overactivation of BMP-signaling results in a ventralized phenotype (Schmidt and Starck, 2004).

The key gradient for left-right patterning in tail morphogenesis is the nodal/spaw gradient, which is initiated by Kupffer’s vesicle (KV), a fluid-filled transient structure localized at the ventral posterior side of the embryo (Kupffer, 1868; Long et al., 2003). Cilia within Kupffer’s vesicle generate an anti-clockwise fluid flow, which in turn induces asymmetric expression of signaling genes such as spaw and establish a nodal/spaw gradient (Essner et al., 2005). This is not only essential for left-right patterning of the tail, but also for asymmetrical organ placement elsewhere in the embryo. Impairment of this signaling pathway can be induced by various drugs (Levin, 2005) and causes tail defects, randomization of organ positioning and heart looping failure (Bonetti et al., 2014; Kramer-Zucker et al., 2005; Smith et al., 2011).

Finally, anterior-posterior patterning is controlled by antagonistic gradients of Wnt/Fgf and retinoic acid, which regulate each other through feedback mechanisms (Stulberg et al., 2012). Key genes in this process are *cyp26a1* and *aldh1a2*. *Cyp26a1* is expressed in the presomitic mesoderm of the tail bud, and degrades retinoic acid into retinoid metabolites (Chithalen et al., 2002; Kudoh et al., 2001; Roberts, 2020). *Aldh1a2* is expressed more anteriorly and produces retinoic acid (Pittlik and Begemann, 2012). High levels of retinoic acid inhibit Wnt/Fgf signaling and conversely, expression of *aldh1a2* is inhibited by high levels of *fgf8a* (Draut et al., 2019). Furthermore, retinoic acid signaling regulates body axis extension and somitogenesis by controlling expression of downstream homeobox transcription factor genes, including *hox* and *cdx* (Shimizu et al., 2006). Fgf/Wnt signaling controls transcription factors such as *msgn1* and *tbx16*, which in turn regulate neuromesodermal stem cells that reside in the tail bud (Manning and Kimelman, 2015). Perturbations of retinoic acid signaling in zebrafish result in severe tail defects. For example, mutants of *cyp26a1* (also known as *gir*) show a shortened and distorted tail (Emoto et al., 2005). Incubation of zebrafish embryos with retinoids causes severe pleiotropic defects including truncation and shortened body axis (D’Aniello et al., 2015; Minucci et al., 1996).

Taken together, morphogen gradients regulate signaling pathways in an intricate manner and shape the developing embryo. Modulating these signaling pathways may particularly affect tail formation. Small molecules that affect zebrafish tail morphogenesis may provide new insights into morphogenesis on the one hand and into signaling pathways on the other.

Here, we found ectopic tail formation in response to a fungal filtrate. We established that monoterpene alcohols, in particular geraniol, induced ectopic tail formation during a 4h window at tail bud stage, most potently in Tüpfel long fin (TL) embryos. We show that the mode-of-action of geraniol is distinct from known ectopic tail inducing compounds such as BMP-pathway inhibitors and para- coumaric acid methyl ester (Gebruers et al., 2013; Yang and Thorpe, 2011). Using transcriptomics of tail buds at tail bud stage, we found genes involved in retinoic acid signaling to be downregulated in geraniol treated embryos. Yet, overexpression of these genes or co-incubation with enhancers of retinoic acid signaling did not rescue the phenotype. Next, time-lapse imaging revealed a migration defect of Kupffer’s vesicle derived cells (KVDCs), suggesting that impaired migration of these cells underlies geraniol-induced ectopic tail formation. Finally, when looking at later stages, we found expression of presomitic mesoderm, somite and notochord markers in ectopic tails, but no neural markers. Conversely, all these markers were detected in the main tail, but expression of Kupffer’s vesicle markers was lacking following geraniol treatment. We conclude that former Kupffer’s vesicle epithelial cells function as a distinct cluster of tail progenitors and their migration is essential for normal tail morphogenesis.

## Results

### Monoterpenes cause ectopic tail formation in zebrafish embryos

Previously, we reported a phenotypic screen of 10,207 fungal filtrates on developing zebrafish embryos (Hoeksma et al., 2019). The filtrate of the fungus *Ceratocystis populicola* (CBS 119.78) was found to induce ectopic tail and fin formation on the ventral side of zebrafish embryos, accompanied by a split notochord and a shorter body axis compared to untreated embryos at 48 hours post fertilization (hpf) (Fig. 1A,B). We selected this fungus for purification and identification of the active compound(s) and to examine the phenotype further.

**Fig. 1.**
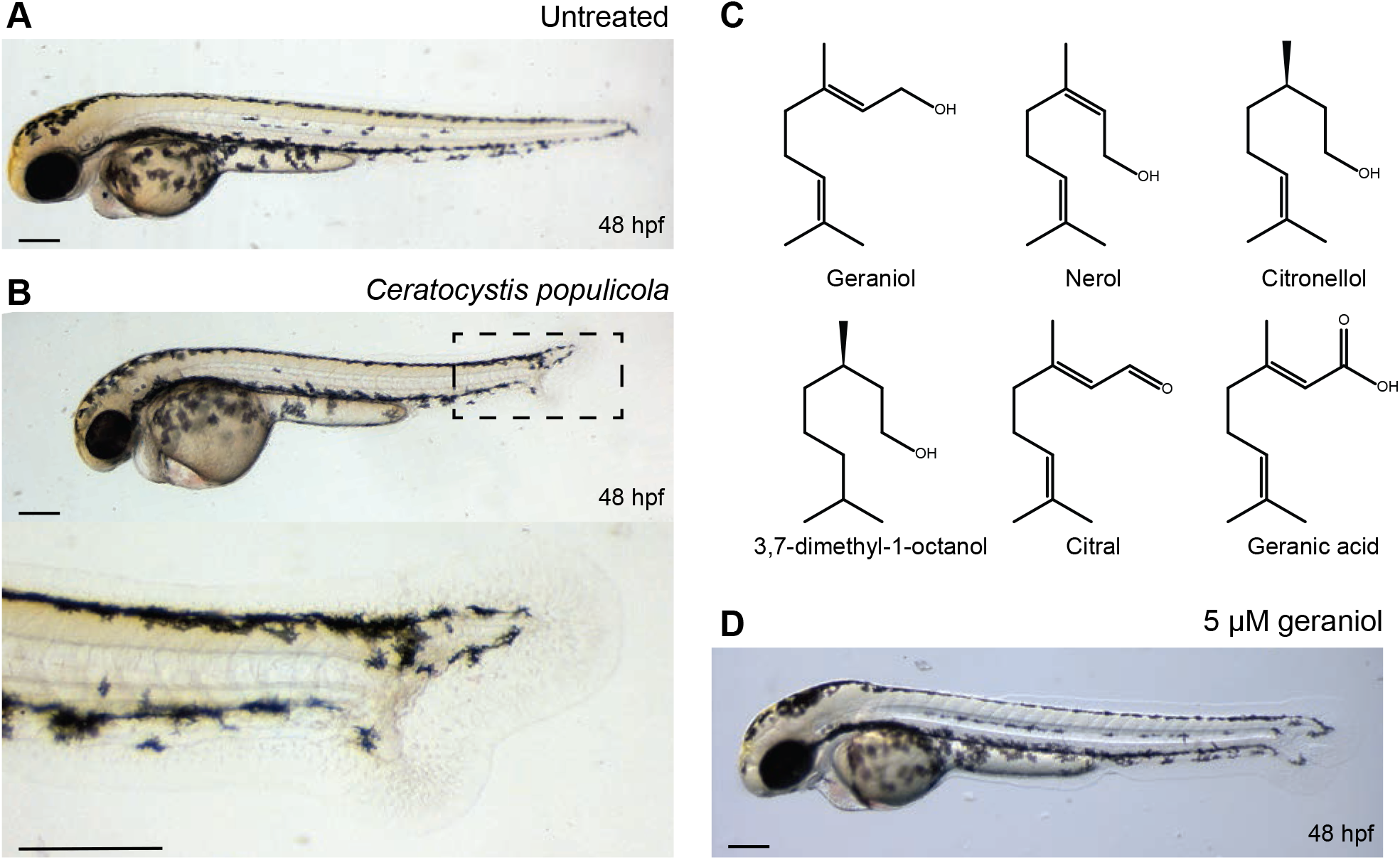
Monoterpenes from *C. populicola* induce an ectopic tail. (A) Untreated control (48 hpf). (B) An example of ectopic tail phenotype induced by the filtrate of C*. populicola*, treatment 6-48 hpf. (C) Molecular structures of compounds inducing ectopic tail. (D) An example of phenotype induced by 5 µM geraniol, treatment 8-24 hpf. Scale bar = 200 µm.

Since Ceratocystis species are known to produce a wide range of volatile compounds (Lanza and Palmer, 1977; Sanchez et al., 2002; Sprecher and Hanssen, 1983), we attempted to purify the active compound through distillation (Fig S1A) and tested the resulting fractions on zebrafish embryos. The most potent fraction was analyzed first by LC-MS revealing a distinct fragmentation pattern specific for the active compound at m/z = 137.2, 95.1 and 81.1 (Fig. S1B). High resolution mass spectrometry revealed a monoisotopic mass of 177.1255 (M+Na^+^), resulting in a likely molecular formula of C_10_H_18_O (177.1256, calculated for C_10_H_18_ONa^+^).

Various monoterpene alcohols match this molecular formula of C_10_H_18_O and distinct mass spectrometry fragmentation pattern (Holzinger et al., 2000; Steeghs et al., 2007). Moreover, monoterpene alcohols are known to be produced by Ceratocystis species (Sprecher and Hanssen, 1983). Therefore, we obtained a selection of commercially available monoterpene alcohols (Fig. 1C) and tested them on zebrafish embryos in serial dilutions for their ability to induce ectopic tails (Fig. S2). Of the monoterpene alcohols, geraniol showed the best capacity to induce ectopic tails at concentrations as low as 2 µM (Fig. S2A). Throughout this paper we used 5 µM, because it reliably induced ectopic tails (Fig 1D). Nerol, a stereoisomer of geraniol, was slightly less potent with a minimal inducing concentration (MIC) of 5 µM (Fig. S2B). In contrast, linalool and cyclic monoterpene alcohols such as eucalyptol and α-terpineol were not able to induce ectopic tails up until a concentration of 500 µM. In addition, we also tested closely related monoterpenes, such as geranic acid (C_10_H_16_O_2_), the carboxylated form of geraniol, which was able to induce ectopic tails at a lowest concentration of 1 µM (Fig. S2C).

Other closely related compounds such as 3,7-dimethyl-1-octanol (C_10_H_22_O), citral (C_10_H_16_O) and citronellol (C_10_H_20_O) (Fig. 1C) were also able to induce ectopic tails albeit at 100 µM, 25 µM and 25 µM respectively (Fig. S2D-F). All compounds that induced ectopic tails, induced increasingly severe pleiotropic phenotypes at higher concentrations. In conclusion, we identified monoterpene alcohols, particularly geraniol, as the active compounds from *Ceratocystis populicola* that induced ectopic tail formation in zebrafish embryos, and in the process, we identified chemically related monoterpenes to induce ectopic tails as well.

### Geraniol-induced ectopic tail formation is zebrafish strain dependent

Next to investigate the morphological defects after 48 hpf, we treated embryos with 5 µM geraniol from 8 until 24 hpf and tracked tail development until 5 dpf. Variable tail phenotypes were observed, which were classified in three distinct categories (Fig. 2A): normal tail development, a severe phenotype for tails with clear ectopic tissue and a split notochord, and a mild phenotype, where the notochord is split at the tail tip, but no ectopic tissue is formed. Strikingly, distinct responses to geraniol treatment were observed when using different zebrafish strains. When using Tüpfel long fin (TL) embryos, 94.4% were found to develop a severe phenotype upon geraniol treatment, with the remaining 5.6% developing a mild phenotype (Fig. 2B). Overall, treated TL embryos were on average 15.3% shorter than untreated embryos (Fig. 2C). In contrast, 91,2% of geraniol-treated WIK embryos showed normal tail embryos, no embryos exhibited the severe phenotype and only 8,8% displayed a mild phenotype. Overall, treated WIK embryos were slightly shorter (3.7%) than untreated embryos. While higher concentrations of geraniol did not increase the number of ectopic tails in WIK embryos, a pleiotropic phenotype, similar to the phenotype observed in TL embryos, was induced. Finally, treatment of embryos from crosses of TL with WIK resulted in fewer ectopic tails (72.2%) and on average the embryos were 11.9% shorter.

**Fig. 2.**
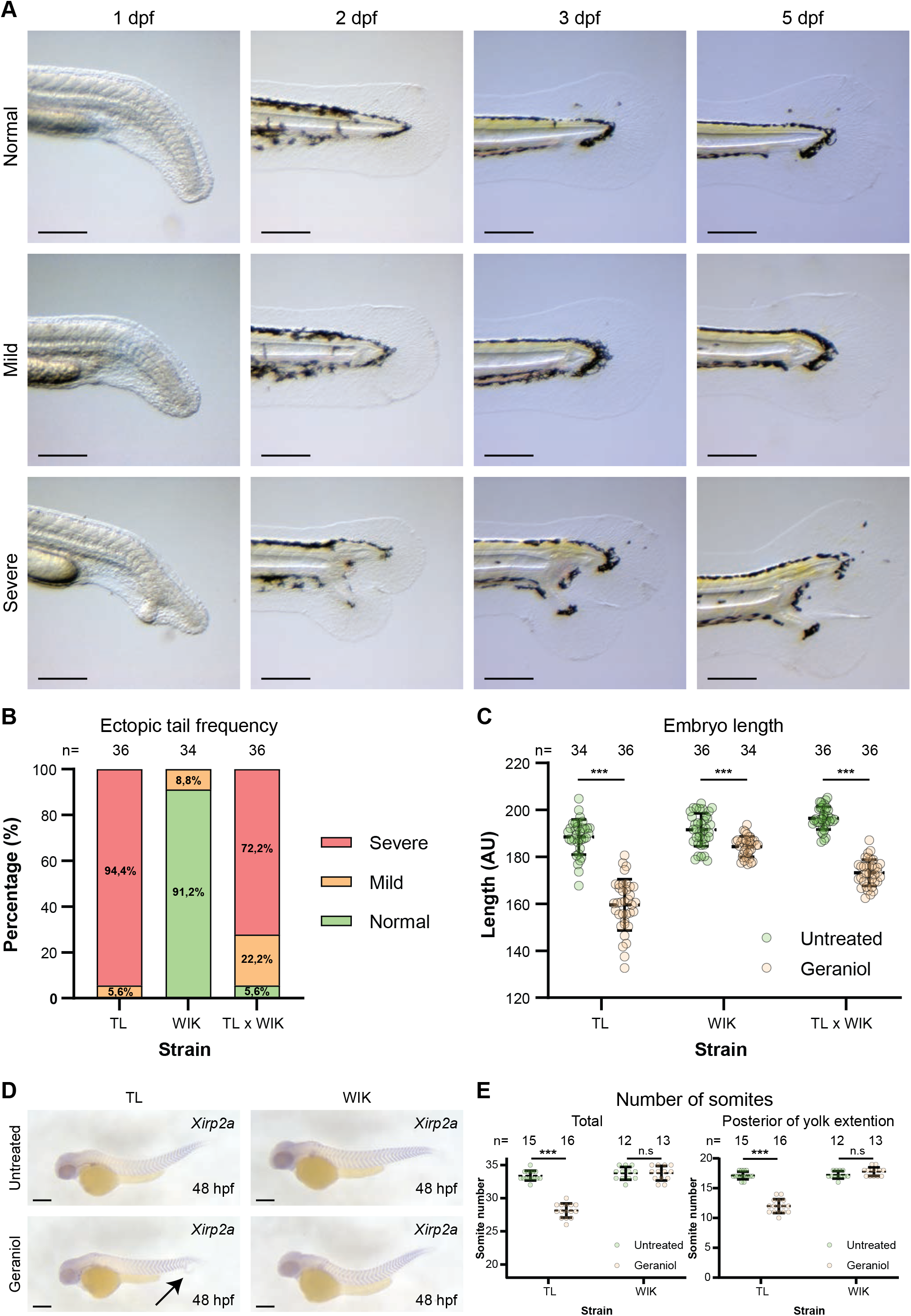
Geraniol induced an ectopic tail in TL, but not WIK embryos. (A) Variations of phenotypes caused by 5 µM geraniol (treatment 8-24 hpf). Representative examples of a normal, a mild and a severe embryo, imaged at the indicated times as indicated, are shown. (B) Phenotype frequency per zebrafish strain using TL and WIK family crosses and single crosses of TL with WIK. Total number of embryos (n) is indicated for each strain. (C) Quantification of zebrafish embryo length in arbitrary units at 48 hpf per zebrafish strain, untreated and 5 µM geraniol treated as indicated. Results are expressed as mean ± standard deviation. Significance was determined using an unpaired t-test (*** = p< 0.001). Total number of embryos (n) per condition is indicated. (D) Expression pattern of *xirp2a* in untreated and geraniol treated (5 µM, 8-24 hpf) TL and WIK embryos, fixed at 48 hpf was established by *in situ* hybridization and imaged. Representative embryos are shown. (E) The total number of somites was counted as well as the number of somites posterior to the yolk extension. Average number of somites is shown ± standard deviation. Significance was determined using an unpaired t-test (*** = p< 0.001). Scale bar = 200 µm.

To investigate the differences in embryo length upon geraniol treatment in more detail, *in situ* hybridization was performed using a *xirp2a* probe (Fig. 2D), which marks the somite boundaries. Geraniol-treated TL embryos, but not WIK embryos, developed significantly less somites than untreated embryos (Fig 2E). This difference was caused by the reduced number of somites posterior to the yolk extension. Furthermore, in both TL and WIK embryos treated with geraniol, the width of the somites was reduced posterior to the yolk extension compared to untreated embryos, whereas there were no differences in the width of anterior somites. In addition, *xirp2a* expression was observed in ectopic tails, indicating that somites are formed in ectopic tails. In conclusion, geraniol treatment induced ectopic tails and affected somite formation in a zebrafish strain dependent manner.

### Ectopic tail formation initiates at tail bud stage

To determine at which developmental stages geraniol induced ectopic tail formation, we tested geraniol treatment on TL zebrafish embryos for various incubation periods (Fig. 3A). Treatments starting at or before 14 hpf induced ectopic tails in all embryos, whereas starting at either 15 or 16 hpf only induced ectopic tails in 60% and 40% of the embryos, respectively. Treatment starting at 17 hpf or later did not induce any ectopic tails. Conversely, treatment until 18 hpf or later induced ectopic tails in all embryos, whereas treatment until 15, 16 or 17 hpf only induced ectopic tails in part of the embryos and aborting treatment at earlier time points did not induce any ectopic tails. Taken together, incubating embryos with geraniol between 14 and 18 hpf was necessary and sufficient for development of an ectopic tail. This period includes the tail bud stage at approximately 16 hpf (Kimmel et al., 1995).

**Fig. 3.**
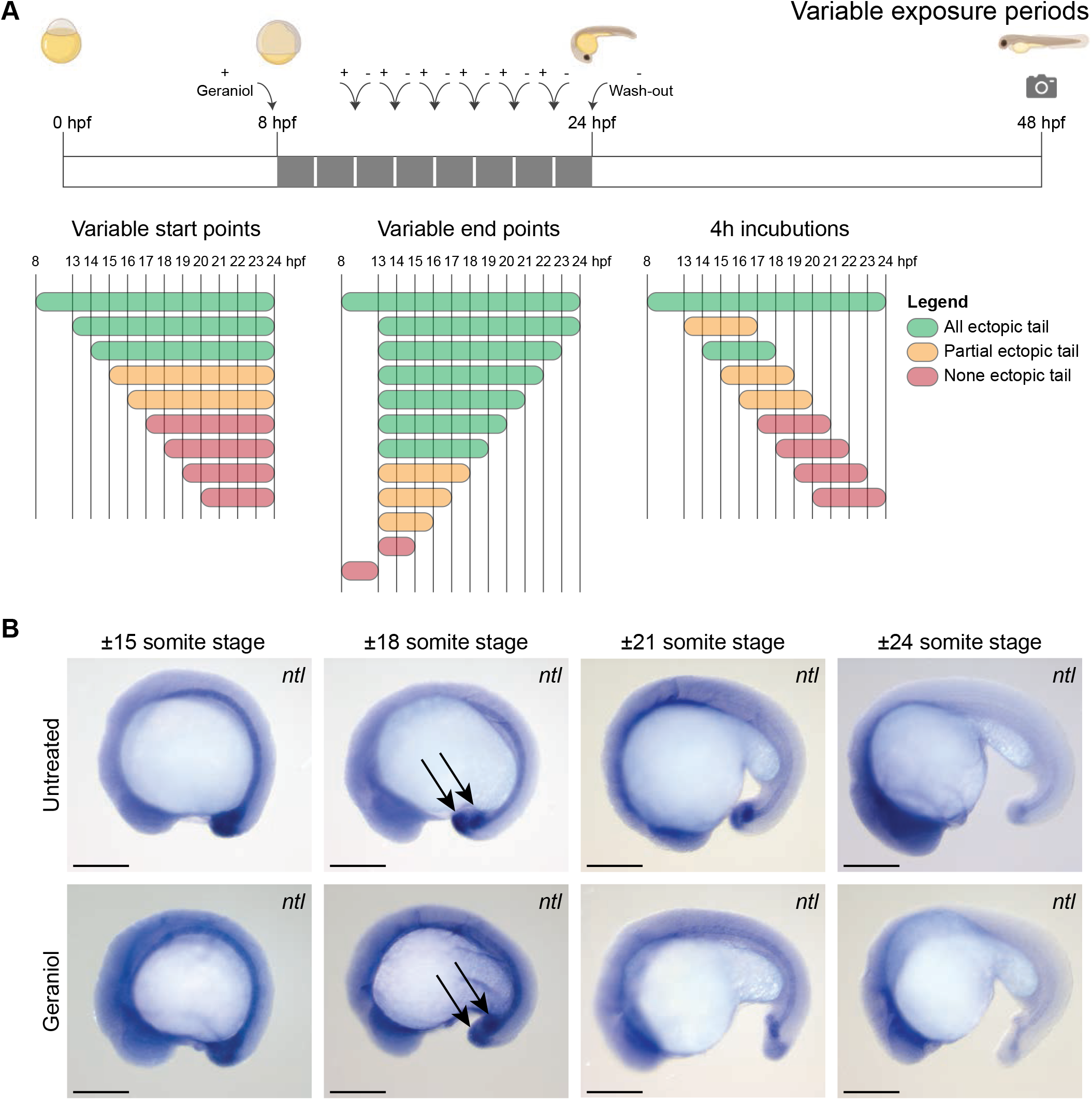
Embryos are sensitive to geraniol between 14 and 18 hpf and geraniol treatment affects *ntl* expression. (A) Embryos treated with 5 µM geraniol for variable incubation periods (5 embryos per condition). Embryos were washed twice with E3 after treatment. Phenotypes were scored at 48 hpf. Green indicates all embryos developed an ectopic tail, orange indicates 1-4 embryos developed an ectopic tail, red indicates no embryos developed an ectopic tail. (B) *In situ* hybridization showing *ntl* expression at 15- to 24- somite stages as indicated. Control (untreated) and embryos treated with 5 µM geraniol from 8 hpf onwards are shown. Arrows highlight the two *ntl* expressing buds at the 18-somite stage in both untreated and geraniol treated embryos. Scale bar = 200 µm.

Next, the morphological events underlying the onset of the phenotype were investigated.

Geraniol treated and untreated embryos were fixed at various time points between approximately 15- somite stage and 24-somite stage. *In situ* hybridization targeting the tail bud and notochord marker *ntl* (also known as *tbxta*) showed no apparent difference at 15- and 18-somite stage between treated and untreated embryos (Fig. 3B). Notably, at 18-somite stage *ntl* appeared to be expressed in two distinct buds in both treated and untreated embryos, with the anterior bud co-localizing with Kupffer’s vesicle. It is noteworthy that *ntl* plays an essential role in maintaining neuromesodermal progenitors (Martin and Kimelman, 2008). The observation that *ntl* was expressed in two distinct buds even in untreated embryos might indicate that there are two separate populations of progenitor cells that give rise to tail. At 21- and 24-somite stage a single bud remained in untreated embryos, whereas in treated embryos the two distinct buds persisted, and a split notochord was observed. This suggests that failure to merge the two *ntl*-positive pools resulted in ectopic tail formation in geraniol-treated embryos.

### Geraniol uses a distinct mechanism compared to other known ectopic tail inducing compounds

Previously, several other small molecules, such as para-coumaric acid methyl ester (pCAME) and BMP-inhibitors dorsomorphin, DMH-1 and cercosporamide were reported to induce ectopic tail tissue (Gebruers et al., 2013; Hoeksma et al., 2020; Yang and Thorpe, 2011). The effect of these compounds was compared by testing them in parallel to geraniol in our assay (Fig. S3) and distinct defects were found. TL-embryos treated with DMH-1 showed a dorsalized phenotype lacking parts of the ventral fin, a phenotype we never observed in geraniol treated embryos, with only a small percentage of the embryos developing ectopic tissue (Fig. S3A). Using pCAME we only observed ectopic tails when performing 1h pulse treatments at 100 µM, similarly to reported (Gebruers et al., 2013), but not during continuous incubation between 8 and 24 hpf (Fig. S3B). Furthermore, to establish whether there is any synergistic effect between these compounds and geraniol, we performed combination treatments using half the MIC and/or the full MIC of the compounds. Treatment with a combination of geraniol and DMH-1 each at full MICs, 2 µM and 250 nM respectively, resulted in a more severe phenotype, but no increase in the number of ectopic tails (Fig. S4A). Also no synergy was observed nor additive defects using half the MIC for geraniol and pCAME (Fig. S4B).

WIK embryos were less sensitive to geraniol. Yet, incubation of WIK embryos with BMP-inhibitor DMH-1 induced loss of ventral fins with a similar frequency as TL-embryos (Fig. S5), indicating that WIK embryos were sensitive to DMH-1. Taken together, geraniol appears to induce ectopic tail formation through a different mechanism than DMH-1 and pCAME.

### Key regulators of retinoic acid signaling are downregulated in tail buds of geraniol treated embryos

To uncover the molecular mechanisms behind the onset of the ectopic tail, whole transcriptome mRNA sequencing was performed on individual tail buds from geraniol-treated or untreated 15-somite stage embryos. Prior to differential gene analysis, the sequencing depth of each sample was checked by determining the number of reads per gene and total read number (Fig. S6A). We excluded 3 samples from differential gene analysis due to low quality.

We then performed differential gene analysis and found 13 significantly downregulated and 4 significantly upregulated genes in the tail bud of geraniol treated embryos compared to untreated (Fig. 4A, S6B). Notable downregulated genes included *cyp26a1*, *fgf8a*, *raraa* and several hox genes, *hoxa13b*, *hoxd12a* and *hoxc11a*, which are all known for their involvement in retinoic acid signaling (Shimizu et al., 2006; Ye and Kimelman, 2020). More specifically, *cyp26a1* encodes an enzyme that metabolizes retinoic acid, *fgf8a* encodes a FGFR ligand, and its expression is regulated by retinoic acid signaling (Hamade et al., 2006). *Raraa* encodes Retinoic Acid Receptor Aa, a retinoic acid-responsive transcription factor.

**Fig. 4.**
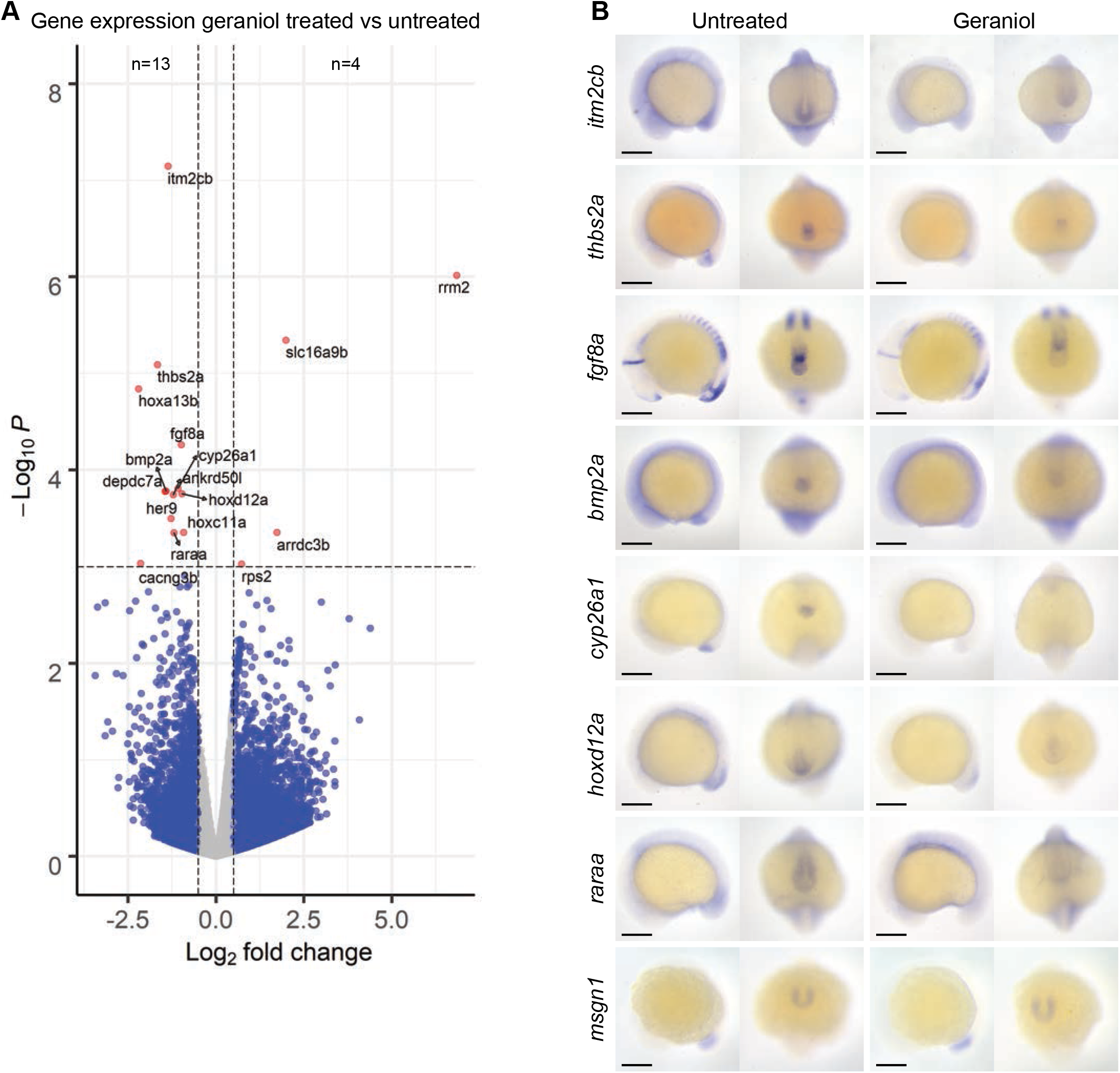
Differential gene expression in response to geraniol treatment. (A) RNA was extracted from the tip of the tail of 15-somite stage embryos untreated (n=7) or treated with 5 µM geraniol from 8 hpf onwards (n=6) and processed for bulk RNA sequencing. Volcano plot shows differentially expressed genes (Total genes included=23,432) between geraniol treated and control embryos. Genes with a fold change of at least ±0.5 and a p-value lower than 0.001 were scored as positive. (B) *In situ* hybridization of selected differentially expressed genes and *msgn1*, imaged laterally and posteriorly. Representative untreated and 5 µM geraniol treated (from 8 hpf onwards) embryos are shown. Scale bar = 200 µm.

Expression of *hoxa13b*, *hoxd12a* and *hoxc11a* is directly controlled by retinoic acid signaling. Moreover, *hoxa13b* is known for its role in mesoderm formation and axis elongation (Ye and Kimelman, 2020). Interestingly, the two most significantly downregulated genes, *itm2cb* and *thbs2a*, have not been associated with zebrafish tail development. Next, the localization of differential gene expression was assessed by *in situ* hybridization using 15-somite stage embryos (Fig. 4B). In agreement with the RNA-sequencing results, lower staining intensity was observed for most of the downregulated genes.

Furthermore, *itm2cb* and *thbs2a* were specifically expressed in the tail bud and Kupffer’s vesicle respectively, which implicates their involvement in tail development, which has not been reported before. We used *msgn1*, a regulator of paraxial mesoderm differentiation of neuromesodermal stem cells, as a control (Manning and Kimelman, 2015; Thisse and Thisse, 2004). *Msgn1* is highly expressed in the tail bud and did not show differential expression in either RNA-sequencing or ISH (Fig. 4B).

*In situ* hybridization using probes for these differentially expressed genes in embryos from WIK incrosses did not show clear differences in expression, whereas quantitative real-time PCR only showed slightly decreased expression of *fgf8a* and increased expression of *itm2cb* between geraniol treated and untreated embryos (Fig. S7). This underlines the strain-specific effect of geraniol and suggests that in TL- embryos, geraniol treatment affects expression of these genes which in turn induces ectopic tail formation.

Differentially down-regulated genes were implicated in retinoic acid signaling. Geraniol has chemical resemblance with retinol with its polyene hydrophobic tail. Interestingly, citral, the aldehyde form of geraniol, has been found to interfere with retinoic acid signaling in *Xenopus* (Schuh et al., 1993). Therefore, we interrogated the potential role of retinoic acid and retinol in ectopic tail formation.

Consistent with previous reports, retinoic acid (50 nM) and retinol (5 µM) induce bent tails (Fig. S8A), but not an ectopic tail, as observed with geraniol (D’Aniello et al., 2015; Minucci et al., 1996). Co-incubation of retinol or retinoic acid with geraniol also did not rescue ectopic tail formation (Fig. S8A). Ectopic tails were still established in addition of the retinoid induced phenotype. Furthermore, *in situ* hybridization (Fig. S8B) and real-time PCR (Fig. S7C,D) indicated upregulation of *cyp26a1* in response to retinoic acid and retinol treatment, contrary to geraniol treatment, whereas *hoxd12a* was severely down regulated in response to treatment with retinoic acid or retinol (Fig. S8C,D). Co-incubation of geraniol with either retinoic acid or retinol still resulted in elevated *cyp26a1* levels compared to control conditions. To investigate a potential causal role, *cyp26a1* was overexpressed by micro-injection of synthetic mRNA encoding *cyp26a1* at the 1-cell stage. Ectopic expression of *cyp26a1* did not rescue geraniol treatment, suggesting that reduced *cyp26a1* expression did not cause ectopic tail formation. Although factors of the retinoic acid signaling pathway were downregulated in response to geraniol, altered retinoic acid signaling does not induce ectopic tail formation.

### Left-right asymmetry is not affected by geraniol treatment

In order to determine the cause of ectopic tail formation, the emergence of ectopic tails in geraniol-treated embryos was studied. Ectopic tails initiate at the site where Kupffer’s vesicle normally collapses and disappears. Kupffer’s vesicle is a transient liquid filled spheroid structure located on the ventral and posterior side of the zebrafish embryo, and plays a crucial role in the establishment of left-right asymmetry in the embryo. To see if geraniol affects left-right asymmetry *in situ* hybridization was performed using the cardiomyocyte marker *myl7* and the liver marker *fabp10a*. No left-right asymmetry defects were apparent in the heart or the liver in response to geraniol treatment (Fig. S9). Whereas there was no difference in liver left-right asymmetry, the liver size in geraniol-treated embryos appeared to be reduced at 48 hpf. However, at 5 dpf, there was no difference in liver size detected anymore, suggesting geraniol treatment induced a transient growth delay of the liver. In conclusion, geraniol does not affect Kupffer’s vesicle-mediated left-right asymmetry.

### Epithelial to mesenchymal transition (EMT) is affected in Kupffer’s vesicle epithelial cells upon geraniol treatment

Previously, it has been shown that upon Kupffer’s vesicle collapse, Kupffer’s vesicle epithelial cells, previously named “Kupffer’s vesicle epithelium derived cells” (KVDCs) (Ikeda et al., 2022), undergo epithelial to mesenchymal transition (EMT)(Amack, 2021; Ikeda et al., 2022). To study the involvement of KVDCs in ectopic tail formation, we performed time lapse imaging using two transgenic lines: *Tg(sox17:EGFP)*, which marks endodermal and Kupffer’s vesicle epithelial cells (Sakaguchi et al., 2006), and *Tg(dand5:EGFP)*, which exclusively marks Kupffer’s vesicle epithelial cells (Ikeda et al., 2022).

We imaged embryos every 15 minutes from before the onset of tail formation, at approximately 10-12-somite stage, until approximately 24 hpf. In untreated embryos for both transgenes, GFP-positive Kupffer’s vesicle epithelial cells first converged inwards into the lumen of Kupffer’s vesicle after which Kupffer’s vesicle disintegrated and ceased to exist (Fig. 5A-C/movie M1,2). Directly after the collapse of Kupffer’s vesicle, the KVDCs transitioned from an epithelial state into a migratory state, a process known as epithelial to mesenchymal transition (EMT). Cells started to dissipate from the main cluster of KVDCs as indicated by the arrows in Fig. 5A-C. In *Tg(sox17:EGFP)* embryos this was reflected in a sprouting pattern upon Kupffer’s vesicle-collapse. In addition, the main cluster of GFP-positive cells migrated towards the tip of the tail, which we quantified by measuring the proximal distance of the fluorescent signal to the tip of the tail (Fig. 5D). When treated with geraniol, Kupffer’s vesicle did disintegrate, but following the collapse, the KVDCs failed to undergo EMT, did not disperse and did not migrate to the tail.

**Fig. 5.**
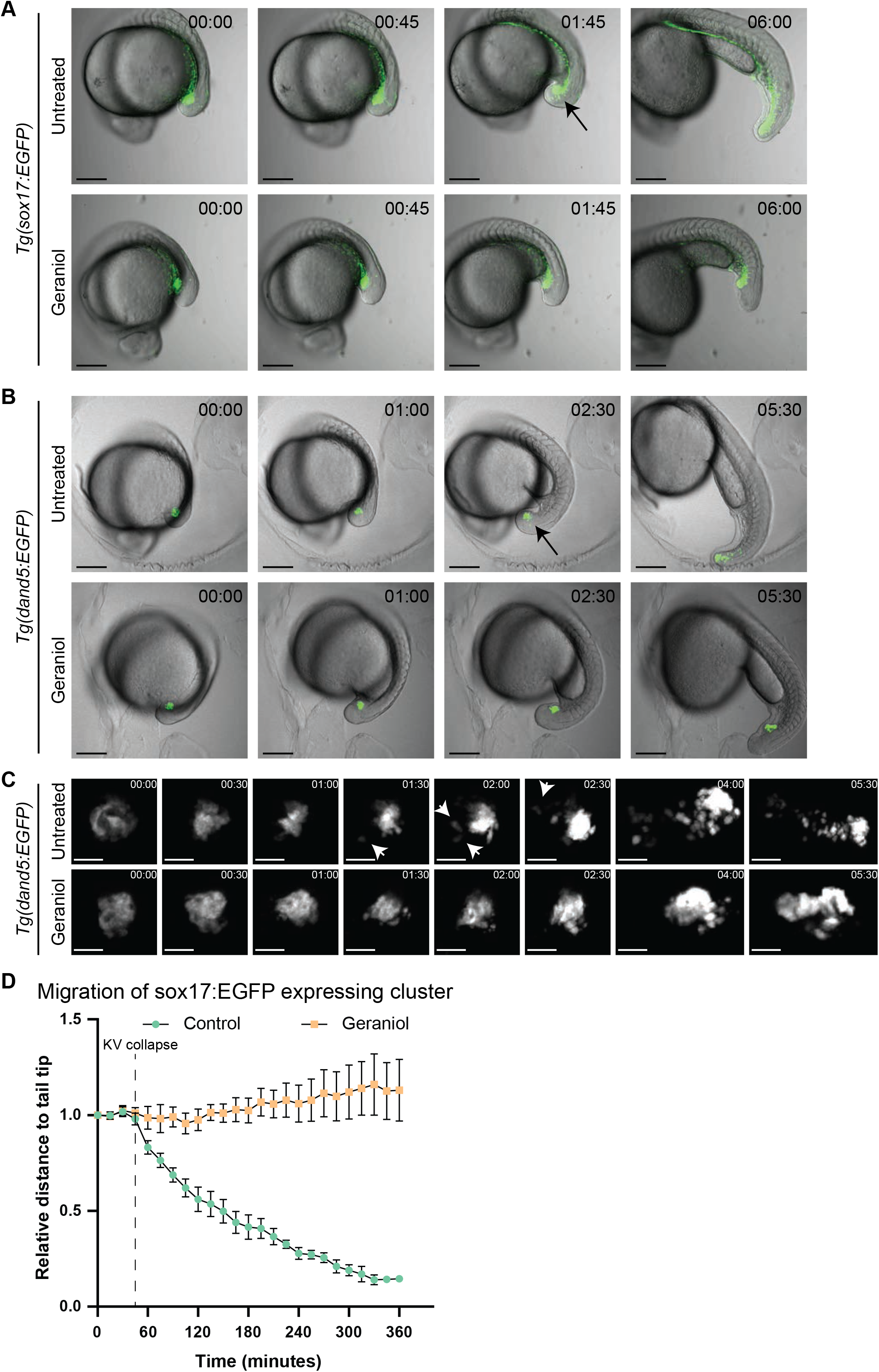
Posterior migration of Kupffer’s vesicle epithelial cells is impaired in response to geraniol treatment. (A) Stills from timelapse imaging (Movie M1) of *Tg(sox17:EGFP)* embryos control (untreated) and treated with geraniol (5 µM from 8 hpf onwards). 00:00 (hh:mm) corresponds to approximately 12- somite stage. (B) Stills from timelapse imaging (Movie M2) using *Tg(dand5:EGFP)* embryos control (untreated) and treated with geraniol (5 µM from 8 hpf onwards). (C) Zoom-in on stills from timelapse imaging using *Tg(dand5:EGFP)* embryos, showing Kupffer’s vesicle collapse in both untreated and geraniol treated embryos (00:00-01:30) and migration of Kupffer’s vesicle epithelial cells. The arrows indicate migratory behavior of fluorescent cells following Kupffer’s vesicle collapse in untreated embryos (01:30-05:30). (D) Quantification of posterior cell cluster migration during timelapse imaging of *Tg(sox17:EGFP)* embryos. Each point in the graph represents the mean distance to the tip of the tail, relative to the distance at the start of the movie. Error bars represent the standard error of the mean. n=5, for both untreated and geraniol treated. Scale bar = 200 µm for figure A and B, 50 µm for figure C.

The proximal distance of *sox17:EGFP*-positive cells to the tip of the tail remained equal throughout tail development (Fig. 5D) and an ectopic tail formed at the former Kupffer’s vesicle site.

Similarly, in untreated *Tg(dand5:EGFP)* embryos cells dispersed from the main cluster after Kupffer’s vesicle-collapse, while the main cluster migrated towards the tail tip (Movie M2/Fig. 5B&C). However, in geraniol treated *Tg(dand5:EGFP)* embryos, the fluorescent cells do not disperse after Kupffer’s vesicle collapse. The fluorescent cells remained restricted to the ectopic tail and did not contribute to the main tail.

Subsequently, expression of these markers was evaluated at 24 hpf and 48 hpf. In untreated embryos, GFP-positive cells from both transgenes were located ventrally of the notochord at 24 hpf and were more dispersed over the posterior end of the tail at 48 hpf. However, in treated *dand5:EGFP* embryos expression was strictly confined to the ectopic tail and likewise, expression of *sox17:EGFP* did not extend into the main tail posterior of the notochord split (fig. 6A). KVDCs do not contribute to the main tail in geraniol treated embryos. Previously, it has been shown that KVDCs contribute to notochord and somite tissue (Ikeda et al., 2022), therefore, we performed *in situ* hybridization to determine which cell types are present in both normal and ectopic tails (Fig. 6B). In geraniol-treated embryos *msgn1*, regulator of paraxial presomitic mesoderm differentiation, was expressed in two distinct clusters, one in the normal and one in the ectopic tail at 24 hpf. This indicates that two separate clusters of presomitic mesoderm and tail progenitor cells exist in embryos with ectopic tails. In addition, other markers such as notochord marker *ntl*, somite and muscle progenitor marker *myod*, and hypochord marker *col2α* were expressed in the ectopic tail as well as the regular tail (Fig. 6B). However, whereas dorsal *col2α* expression appeared as an uninterrupted line, ventral *col2α* expression was interrupted and appeared to deviate into the ectopic tail in treated embryos. Notably, ectopic tails lacked expression of neural ectoderm marker *ngn1* which is only dorsally expressed in the embryo, suggesting KVDCs either lack the capacity or do not receive essential cues to form neural tissue. In conclusion, the imaging and *in situ* hybridization experiments revealed that geraniol treatment blocked migration of KVDCs towards the tip of the tail. Instead, these cells contributed to an ectopic tail bud, which expresses all markers of the normal tail, except the neural marker, *ngn1*. Under normal conditions - in the absence of geraniol – the KVDCs migrate towards the posterior and contribute to the developing tail.

**Fig. 6.**
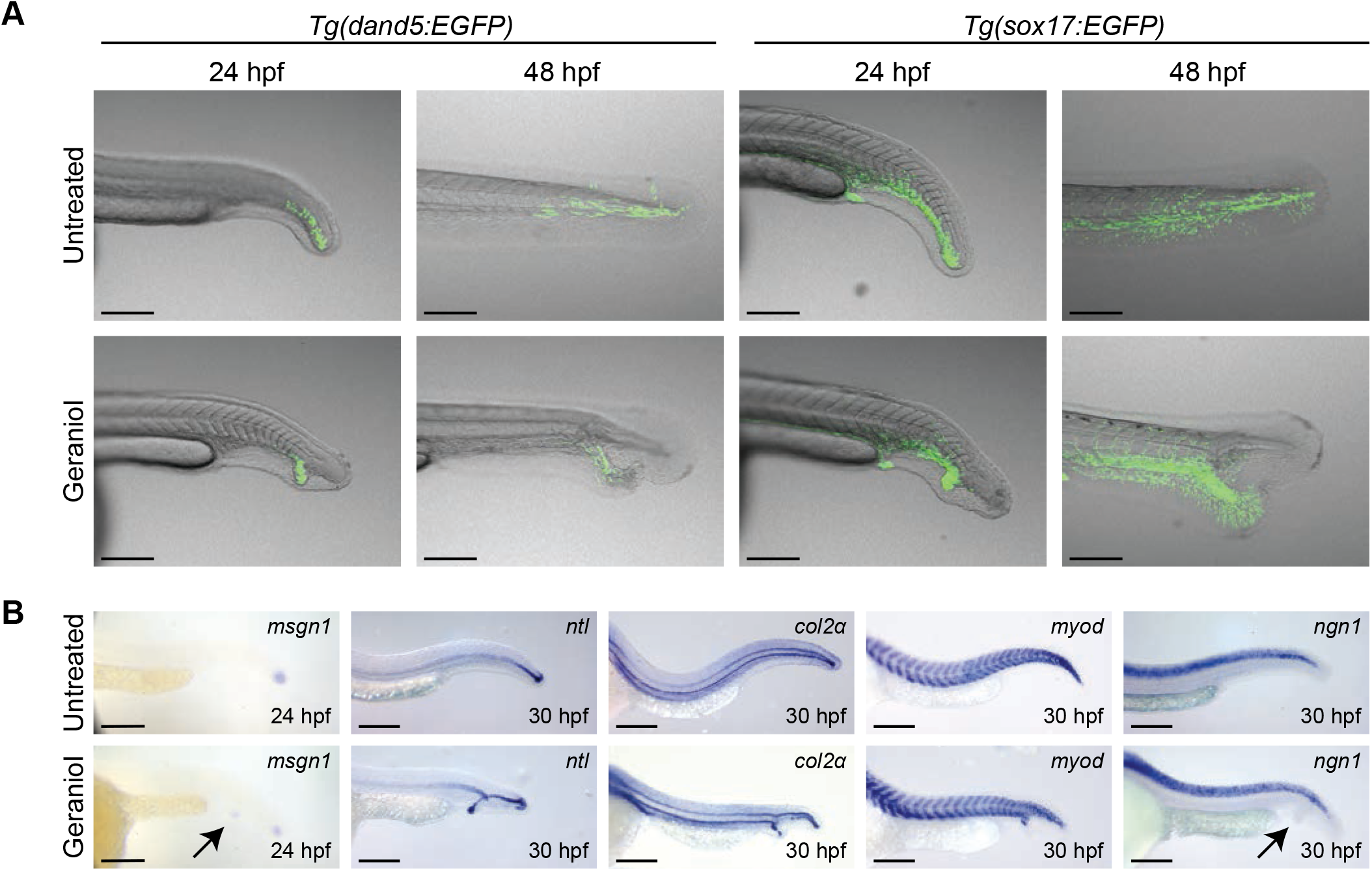
Geraniol treatment caused impaired cell migration and impaired cell fate determination in the ectopic tail. (A) Representative embryos showing expression of *dand5:EGFP* and *sox17:EGFP* at 24 hpf and 48 hpf. Control (untreated) and embryos treated with 5 µM geraniol from 8 – 24 hpf are depicted. (B) Control (untreated) and treated (5 µM geraniol from 8 – 24 hpf) embryos were fixed at 24 hpf for *msgn1,* and 30 hpf and processed for *in situ* hybridization using various markers. Scale bar = 200 µm.

## Discussion

Here, we showed that monoterpene alcohols, in particular geraniol, from the fungus *Ceratocystis populicola* induced ectopic tail formation in TL zebrafish embryos during tail bud stage. Treatment with geraniol affected retinoic acid signaling, however the precise role of this pathway in ectopic tail formation is not clear. Finally, we showed that migration of KVDCs is affected by geraniol, resulting in prolonged separation of two progenitor pools and eventually ectopic tail formation.

Canonically, neuromesodermal progenitor cells reside in the tailbud from which they first undergo EMT and move into the maturation zone under the influence of *msgn1* and *tbx16*, and become migratory. Subsequently, mesodermal cells continue to move anteriorly through the pre-somitic mesoderm after which they start to form somites and contribute to both mesodermal and neural lineages (Chalamalasetty et al., 2014; Manning and Kimelman, 2015). Our results show that KVDCs act as a separate cluster of progenitor cells with the expression of progenitor markers such as *msgn1* and *ntl*. However, at later stages, the ectopic tail of geraniol-treated embryos contained somite and notochord tissue, but not neural tissue (Fig. 2D & 6B). This suggests that KVDCs can only transdifferentiate into mesodermal lineages, not neural lineages or potentially lack cues to differentiate into other lineages.

The extent of the plasticity of KVDCs is an interesting topic for future research. Nonetheless, the lack of posterior *Tg(dand5:EGFP)* and *Tg(sox17:EGFP)* expression in the regular tail in 24 hpf old geraniol treated embryos (Fig. 6A) and the integration of KVDCs in the ventral-posterior tail under normal conditions demonstrate that KVDC migration is essential for tail morphogenesis.

Geraniol may act on several signaling pathways to drive ectopic tail formation. Previously, ectopic tail formation has mostly been associated with effects on BMP-signaling and non-canonical Wnt-signaling (Gebruers et al., 2013; Yang and Thorpe, 2011). Our results show that geraniol-induced ectopic tail formation was not due to BMP-signaling, because we did not observe any synergistic effects of treatments combining geraniol with DMH-1 or pCAME. In addition, the BMP-signaling inhibitor, DMH-1, had a similar penetrance in embryos from both TL and WIK strains, whereas geraniol only induced ectopic tails in TL embryos.

The difference in effect of geraniol on TL and WIK strains (Fig. 2) may provide an opportunity to deduce the targets or pathways involved in ectopic tail formation by geraniol. Genetic comparison of TL and WIK strains may provide clues regarding the targets of geraniol. Extensive genetic variations exist between wild type strains, which may affect the outcome of experiments, including compound screens (Guryev et al., 2006). It is likely that the target of geraniol is either differentially expressed between WIK and TL strains or harbors single nucleotide polymorphisms (SNPs), which may influence binding or conversion of geraniol. Transcriptomic or genomic approaches could lead to identification of differences between WIK and TL strains and flag potential targets of geraniol.

Based on our RNA expression data (Fig. 4), retinoic acid signaling appeared to be a good candidate pathway for involvement in ectopic tail formation. Furthermore, in WIK embryos we did not observe differential expression of retinoic acid genes. Moreover, citral, the aldehyde form of geraniol, was previously reported to interfere with retinoic acid signaling in *Xenopus* (Schuh et al., 1993). Geraniol has chemical resemblance to retinol with its polyene hydrophobic tail and disruption of the retinoic acid balance in the zebrafish tail was reported to cause an array of tail phenotypes (Rydeen et al., 2015).

However, whereas expression of *cyp26a1* was downregulated by geraniol treatment, it was upregulated by retinoic acid, even when co-treated with geraniol. Upon co-treatment with geraniol and retinoic acid, ectopic tails still formed, suggesting that retinoic acid was not directly involved in ectopic tail formation. Potentially, differential expression of other genes that we identified, has a causal role. For instance, *arrdc3a*/ARRDC3 has recently been reported to inhibit EMT in triple knock-out breast cancer and has tumor suppressing properties (Soung et al., 2019). However, the function of *arrdc3* remains to be determined definitively. The mouse homologue of *thbs2*, *Thbs2*, is reported to be involved in cell adhesion and migration of mesenchymal cells in mouse (Kyriakides et al., 1998) and human *THBS2* is reported as a tumor suppressor (Liu et al., 2023). The mouse homologue *Itm2c*, an integral transmembrane protein, has been reported to function in the brain (Choi et al., 2001) and recently human *ITM2C* has been associated with colorectal cancer (Maurya et al., 2023). It will be interesting to investigate the role of these differentially expressed genes in tail morphogenesis and ectopic tail formation in the future, which may in the process provide insights into their role in disease.

Finally, treatment with geraniol for 4h during tail bud stage was sufficient to induce ectopic tails (Fig 3A). At bud stage, two clusters of tail progenitor cells were apparent, given the bifurcated expression of *ntl* in untreated embryos (Fig. 3B), which was resolved during the next phase. In geraniol- treated embryos this bifurcated expression persisted (Fig. 3B). The time-lapse imaging experiments using *Tg(dand5:EGFP)* and *Tg(sox17:EGFP)* reporter lines revealed inhibited migration of KVDCs in response to geraniol treatment. All results combined, we conclude that Kupffer’s vesicle epithelial cells have transdifferentiation and tail morphogenic potential, and are distinct from tail progenitors residing in the caudal tail bud. Upon Kupffer’s vesicle collapse these KVDCs under normal circumstances become migratory and contribute to the patterning of the tail. Geraniol inhibits migration of KVDCs and thereby induces ectopic tail formation. Overall, our results provide new insights into zebrafish tail morphogenesis and ectopic tail formation, and highlight the importance of KVDC migration.

## Materials and Methods

### Embryo screening assay

For the initial experiments, embryos from family crosses of Tüpfel long fin zebrafish were used. The eggs were first washed with fresh E3-medium, before they were divided over 24-well plates with 10 embryos per well in 1000 µL E3-medium. Samples were added in various concentrations and for various durations as mentioned in text and figure legends. Brightfield imaging was performed using either a Leica M165 FC microscope equipped with a Leica DMC5400 camera or a Leica MZFLIII microscope equipped with a Leica DFC320 camera whilst the embryos were mounted in 2% methyl cellulose. ImageJ was used for size and length measurements.

Tüpfel long fin (ZDB-GENO-990623-2 - catalogue nr. 1174-TL) and WIK zebrafish lines (ZDB-GENO- 010531-2 – catalogue nr. 1171-WIK) were originally obtained from EZIRC (Karlsruhe, Germany).

All procedures involving experimental animals were approved by the local animal experiments committee (Koninklijke Nederlandse Akademie van Wetenschappen-Dierexperimenten commissie) and performed according to local guidelines and policies in compliance with national and European law.

Adult zebrafish were maintained as previously described (Aleström et al., 2019).

### Fungus culture and compound purification and identification

The fungus *Ceratocystis populicola* (CBS 119.78) was grown on cornmeal agar (CMA) at 25 °C for 7 days. Next, the agar with mycelium was cut into cubes of approximately 5x5 mm, which were used to inoculate 20x 100 mL bottles containing 50 mL Czepek dox broth + 0,5% yeast extract. The medium was incubated at 25 °C for 7 days, before filter sterilization using a 0.22 µm Millipore steritop filter.

Approximately 125 mL of the fungal filtrate was used for vacuum distillation using the set-up as shown in Fig. S1A using magnetic steering of the filtrate at 250 rpm and with heating increasing every 40 minutes: 50 °C for 0-40 min; 75 °C for 40-80 min; 150 °C for 80-120 min. Samples were collected every 40 minutes.

Samples were analyzed using a LC-MS and HRMS. HRMS was performed on a µQTOF instrument (Micromass Ltd, Manchester, United Kingdom), which was calibrated using sodium formate, followed by injection of the sample mixed with sodium formate inducing sodium adduct ions of the compound in the process.

### Live imaging

Eggs from either *Tg(sox17:EGFP)* or *Tg(dand5:EGFP)* crosses in a TL background were harvested and grown at 28.5 °C until 8 hpf. Embryos were split over two 6 cm dishes, one dish was incubated with 5 µM geraniol (see Table 1 for chemicals and reagents), whilst the other was left untreated. Subsequently, the embryos were grown overnight at room temperature to delay growth. Fluorescent embryos were selected and placed in a glass bottom dish or glass bottom 24-well plate. *Tg(sox17:EGFP)* embryos were (co-)incubated with MS222/tricane during the experiment, *Tg(dand5:EGFP)* embryos were injected with bungarotoxin messenger-RNA (Addgene catalogue #69542) at the one-cell stage to immobilize them.

**Table 1:** Chemicals & Reagents.

| Reagent | Supplier | Catalogue # |
| --- | --- | --- |
| 3,7-dimethyl-1-octanol | Merck | W239100 |
| 3,7-dimethyl-6-octenamide | Merck | S798355 |
| Anti-Digoxigenin-AP, Fab fragments | Merck | 11093274910 |
| α-Terpineol | Merck | 4899 |
| Citral | Merck | W230308 |
| Citronellol | Merck | W2302901 |
| DIG RNA Labeling Mix | Merck | 11277073910 |
| Dimethyl sulfoxide | Merck | D8418 |
| DMH-1 | Merck | D8946 |
| Dorsomorphin | Merck | P5499 |
| Eucalyptol | Merck | C80601 |
| FastStart Universal SYBR Green Master | Merck | 4913914001 |
| Geranic acid, 98% (sum of isomers) | Fisher | 11433028 |
| Geraniol | Merck | 163333 |
| Linalool | Merck | L2602 |
| Methyl cellulose | Merck | M0387 |
| MS222 | Merck | E10521 |
| Nerol | Merck | 50949 |
| p-Coumaric acid | Merck | C9008 |
| Paraformaldehyde | Merck | 158127 |
| Retinoic acid | Merck | R2625 |
| Retinol | Merck | R7632 |
| TRIzol™ Reagent | Fisher | 15596018 |

Embryos were imaged using a Leica SP8 microscope. Z-stacks were obtained every 15 minutes. Maximum projections and quantifications were created using ImageJ.

### *In situ* hybridization

Embryos were fixed in 4% paraformaldehyde either overnight at 4°C or for 3h at room temperature. In situ hybridization was performed as previously described (Thisse and Thisse, 2008). Digoxigenin-labeled probes were generated from PCR products or plasmids. The sequences of the primers used to generate these PCR products are listed in Table S1. The reverse primers all contain a T7 RNA polymerase promoter site preceded by three bases to allow transcription of the PCR product. In situs were imaged using a Leica M165 FC microscope equipped with a Leica DMC5400 camera and a ring light.

### RNA isolation from tail buds

Eggs from TL-crosses were harvested and grown at 28.5 °C until 8 hpf. Embryos were split over two 6 cm dishes, one dish was incubated with 5 µM geraniol, whilst the other was left untreated. Subsequently, the embryos were grown overnight at room temperature to delay growth. At approximately 15-somite stage the eggs were dechorionated and individual tails were isolated by cutting the tail bud using a BD Microlance 3 (30G x ½” - #304000) needle. The tail buds were transferred to a 1.5 mL Eppendorf tube and immediately snap frozen in liquid nitrogen. RNA-extraction was performed using Trizol reagent according to the manufacture’s protocol using 1/10 of the volumes reported. The pellet was resuspended in 10 µL milli-Q water. The quality of the samples was assessed using a bioanalyzer. In total 8 untreated and 8 geraniol treated samples were submitted for RNA sequencing. Library preparation and RNA sequencing was outsourced to Single Cell Discoveries BV, Utrecht. Differential gene analysis was performed using R-studio using R-library deseq2. Volcano plot has been generated using library EnhancedVolcano. Code used for analysis available at request.

### Quantitative PCR

For quantitative PCR five embryos per condition were pooled and snap frozen in liquid nitrogen. Subsequently, RNA-extraction was performed using Trizol reagent according to the manufacture’s protocol. Quantitative real-time PCR was performed using FastStart Universal SYBR Green Master Mix on a Bio-Rad CFX384 Touch Real-Time PCR Detection System with primers as mentioned in table S2. For each condition 3 biological replicates were used and in turn each gene was measured in either triplicate or quadruplicate. As internal reference housekeeping gene *β-actin* was used. Analysis was performed using the Bio-Rad CFX Maestro software. Plots were generated using Prism GraphPad.

## Acknowledgements

We would like to thank Hiroyuki Takeda and Takafumi Ikeda from University of Tokyo for the *Tg(dand5:EGFP)* reporter line and Jeroen Bakkers and Melanie Fremery for the *Tg(sox17:EGFP)* reporter line. Furthermore, we would like to thank Rob Liskamp, Geert-Jan Boons, John Kruijtzer, Albert Heck and Cees Versluis from Utrecht University for their help with the purification and identification of the monoterpenes, Single Cell Discoveries for their help with RNA sequencing, Hubrecht Imaging Centre for their help with confocal imaging and Marloes Blotenburg for her help with sequence mapping and critical review of the paper.

## Author contributions

Conceived and designed the experiments: JH, JdH. Performed the experiments: JH. Analyzed the data: JH. Wrote the paper: JH, JdH.

## Supplementary Material

Fig. S1. Isolation and identification of ectopic tail inducing compound. (A) Representation of vacuum distillation set-up. (B) Mass spectrogram of distillation fraction 2 compared to a background spectrum reveals monoterpene alcohol specific peaks.

Fig. S2. Concentration-dependent ectopic tail formation by monoterpenes. Phenotypes induced by treatment of zebrafish embryos from 8 hpf onwards with increasing concentrations of monoterpenes, including (A) geraniol (B) nerol (C) geranic acid (D) 3,7-dimethyl-1-octanol (E) citral and (F) citronellol. The lowest concentration for each monoterpene shown here is the highest concentration that did not induce ectopic tail formation. Scale bar = 200 µm.

Fig. S3. Tail defects induced by DMH-1 and PCAME. (A) Phenotypes induced by increasing concentrations of BMP-inhibitor DMH-1, including loss of ventral fins and ectopic tissue formation in some embryos as indicated by the arrows. (B) Phenotypes induced by increasing concentrations of para-coumaric acid methyl ester (pCAME). pCAME only induced ectopic tails when applying 1h pulse treatments between 11-12 hpf. Scale bar = 200 µm.

Fig. S4. Geraniol does not cooperate with DMH-1 and pCAME. (A) Combination treatments of various concentrations of geraniol and DMH-1. (B) Combination treatments of various concentrations of geraniol and pCAME. Scale bar = 200 µm.

Fig. S5. DMH-1 induces a similar phenotype in TL and WIK embryos. (A) Phenotype frequency induced by DMH-1 in TL and WIK embryos. (B) Quantification of zebrafish embryo length in arbitrary units at 48 hpf per zebrafish strain in untreated and DMH-1 treated embryos.

Fig. S6. RNA quality control and RNA expression counts. (A) Reads per gene per sample of the 8 control (untreated) and 8 geraniol-treated (5 µM, from 8 hpf onwards). Samples with a low number of reads (in red) were omitted from the RNA-sequencing results. (B) Normalized gene counts of differentially expressed genes.

Fig. S7. Lack of differential gene expression in response to geraniol treatment in WIK embryos. (A) *In situ* hybridization using WIK embryos and probes specific for selected genes that were differentially expressed in TL embryos (Fig. 4B). Note that no clear reduction in expression was observed upon geraniol treatment. (B) Quantitative real-time PCR on 15-somite stage WIK embryos (5 embryos per sample). Only a small decrease in *fgf8a* expression and an increase in *itm2cb* expression were observed.

Fig. S8. Geraniol treatment does not cooperate or interfere with retinoic acid and/ or retinol treatment of embryos. (A) Tail defects induced by 50 nM retinoic acid or 5 µM retinol (8-24 hpf treatment), imaged at 48 hpf, without and with co-treatment with geraniol (5 µM). (B) *In situ* hybridization of TL embryos treated with retinoic acid or retinol using probes specific for selected genes that were differentially expressed in TL embryos in response to geraniol (Fig. 4B). A clear increase in *cyp26a1* expression was observed. Several embryos showed a skewed expression of *fgf8a* around Kupffer’s vesicle. (C) Quantitative real-time PCR on whole 15-somite stage embryos (5 embryos per sample) untreated of treated with either geraniol, retinoic acid or co-treated with 5 µM geraniol and 50 nM retinoic acid. (D) Quantitative real-time PCR on whole 15-somite stage embryos (5 embryos per sample) untreated of treated with either geraniol, retinoic acid or co-treated with 5 µM geraniol and 5 µM retinol.

Fig. S9. Geraniol does not affect left-right asymmetry. *In situ* hybridization using TL embryos (48 hpf and 5 dpf) that had or had not been treated with geraniol (5 µM, 8-24 hpf) and probes specific for *myl7* (cardiomyocytes) or *fabp10a* (liver). It is evident that left-right asymmetry was not affected by geraniol treatment. However, liver size was reduced at 48 hpf.

