## Supplemental Material for "Migration of Kupffer’s vesicle derived cells is essential for tail morphogenesis in zebrafish embryos"

Fig S1

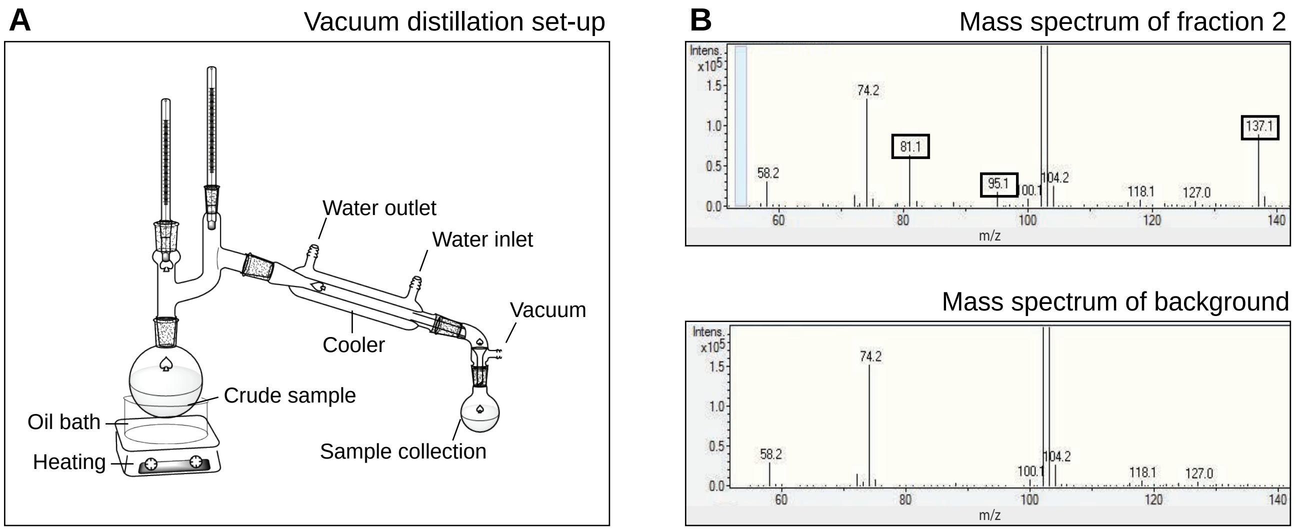
A
Vacuum distillation set-up
Water outlet
Water inlet
Vacuum
Cooler
Crude sample
Oil bath
Heating
Sample collection
B
Mass spectrum of fraction 2
Intens.
x10^5
74.2
81.1
58.2
95.1
100.1
104.2
118.1
127.0
137.1
m/z
Mass spectrum of background
Intens.
x10^5
74.2
58.2
100.1
104.2
118.1
127.0
m/z

Fig S2

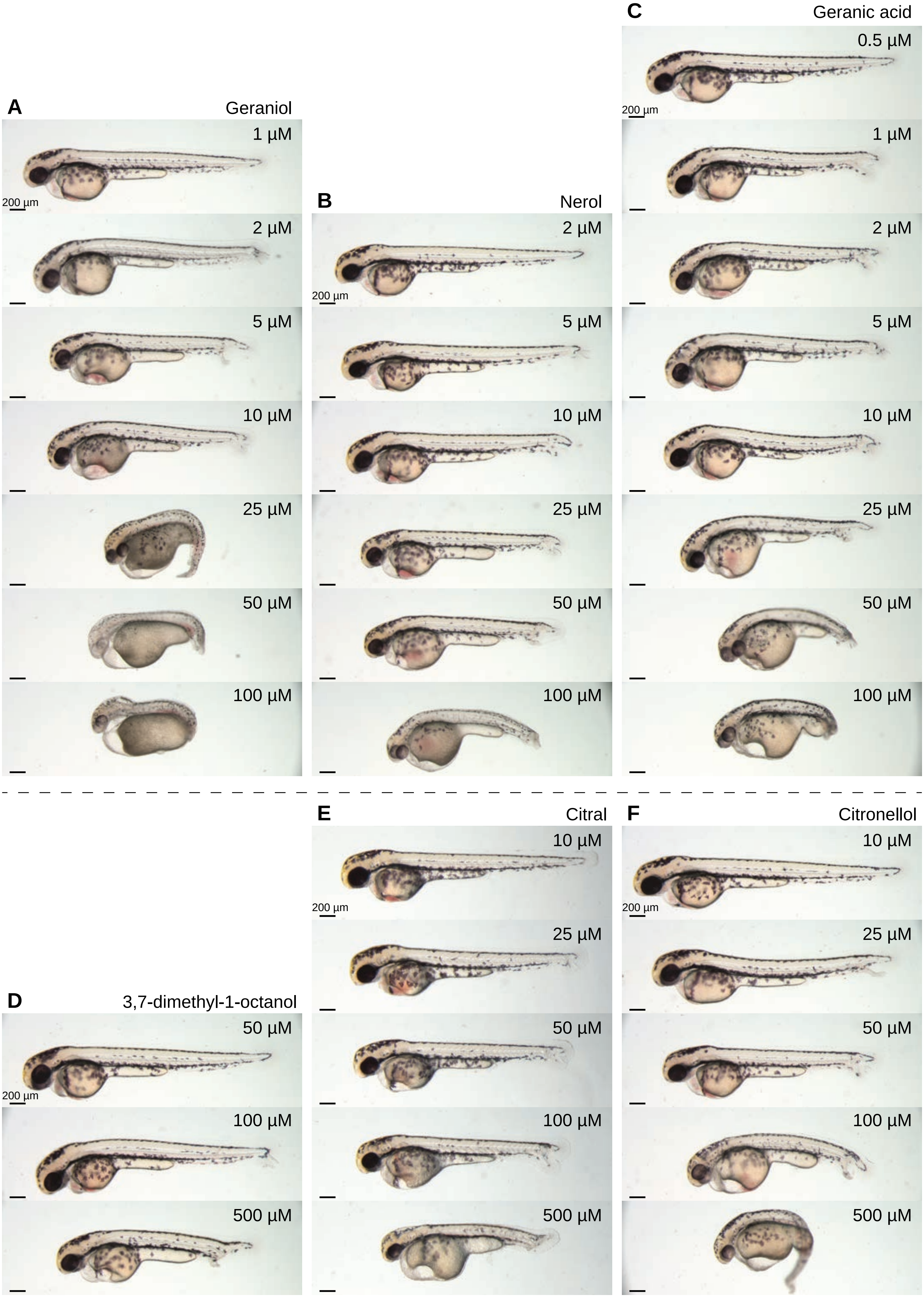

A
Geraniol
1 µM
2 µM
5 µM
10 µM
25 µM
50 µM
100 µM
200 µm
B
Nerol
2 µM
5 µM
10 µM
25 µM
50 µM
100 µM
200 µm
C
Geranic acid
0.5 µM
1 µM
2 µM
5 µM
10 µM
25 µM
50 µM
100 µM
200 µm
D
3,7-dimethyl-1-octanol
50 µM
100 µM
500 µM
200 µm
E
Citral
10 µM
25 µM
50 µM
100 µM
500 µM
200 µm
F
Citronellol
10 µM
25 µM
50 µM
100 µM
500 µM
200 µm

Fig S3

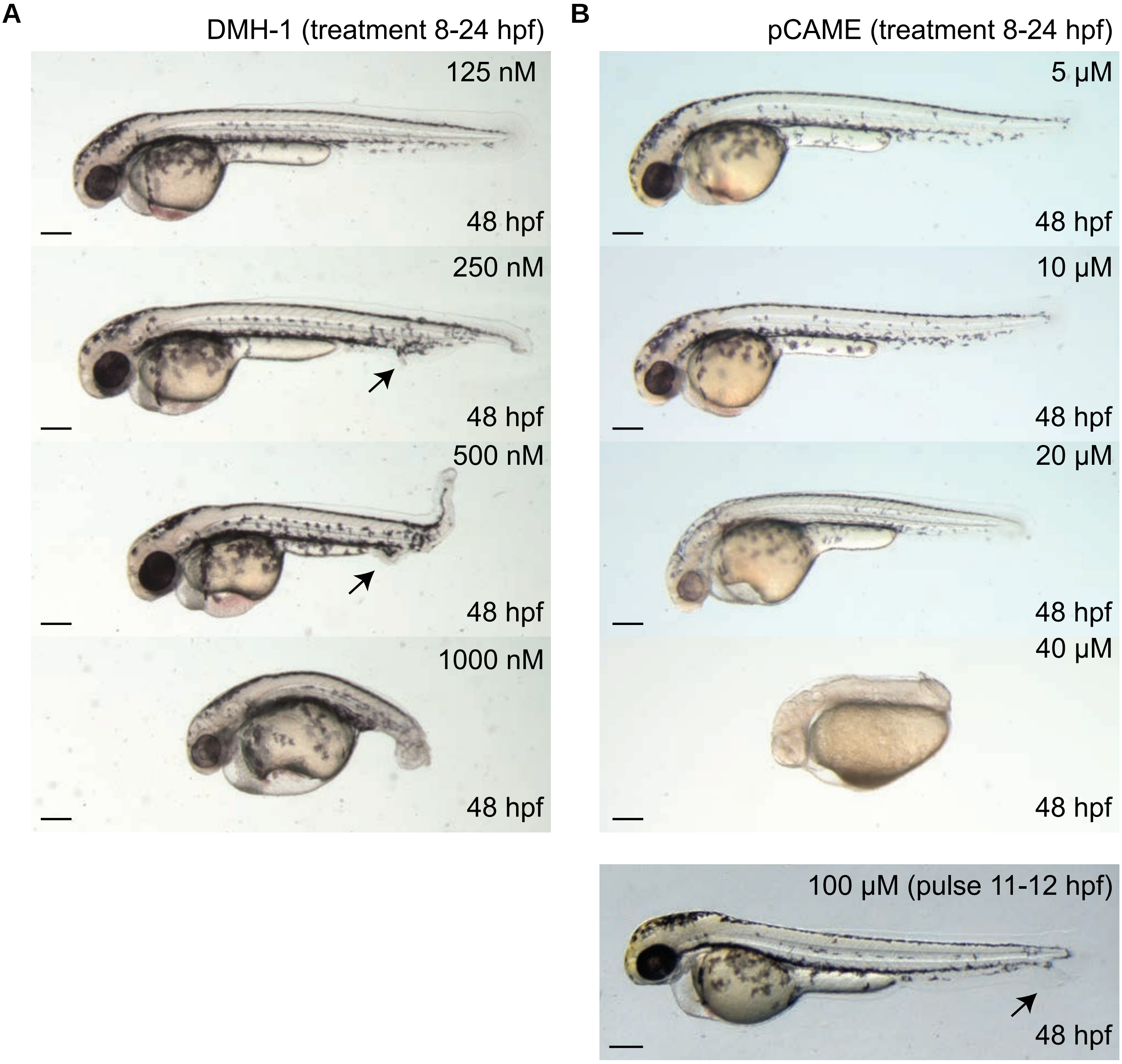
A
DMH-1 (treatment 8-24 hpf)
125 nM
48 hpf
250 nM
48 hpf
500 nM
48 hpf
1000 nM
48 hpf
B
pCAME (treatment 8-24 hpf)
5 µM
48 hpf
10 µM
48 hpf
20 µM
48 hpf
40 µM
48 hpf
100 µM (pulse 11-12 hpf)
48 hpf

Fig S4

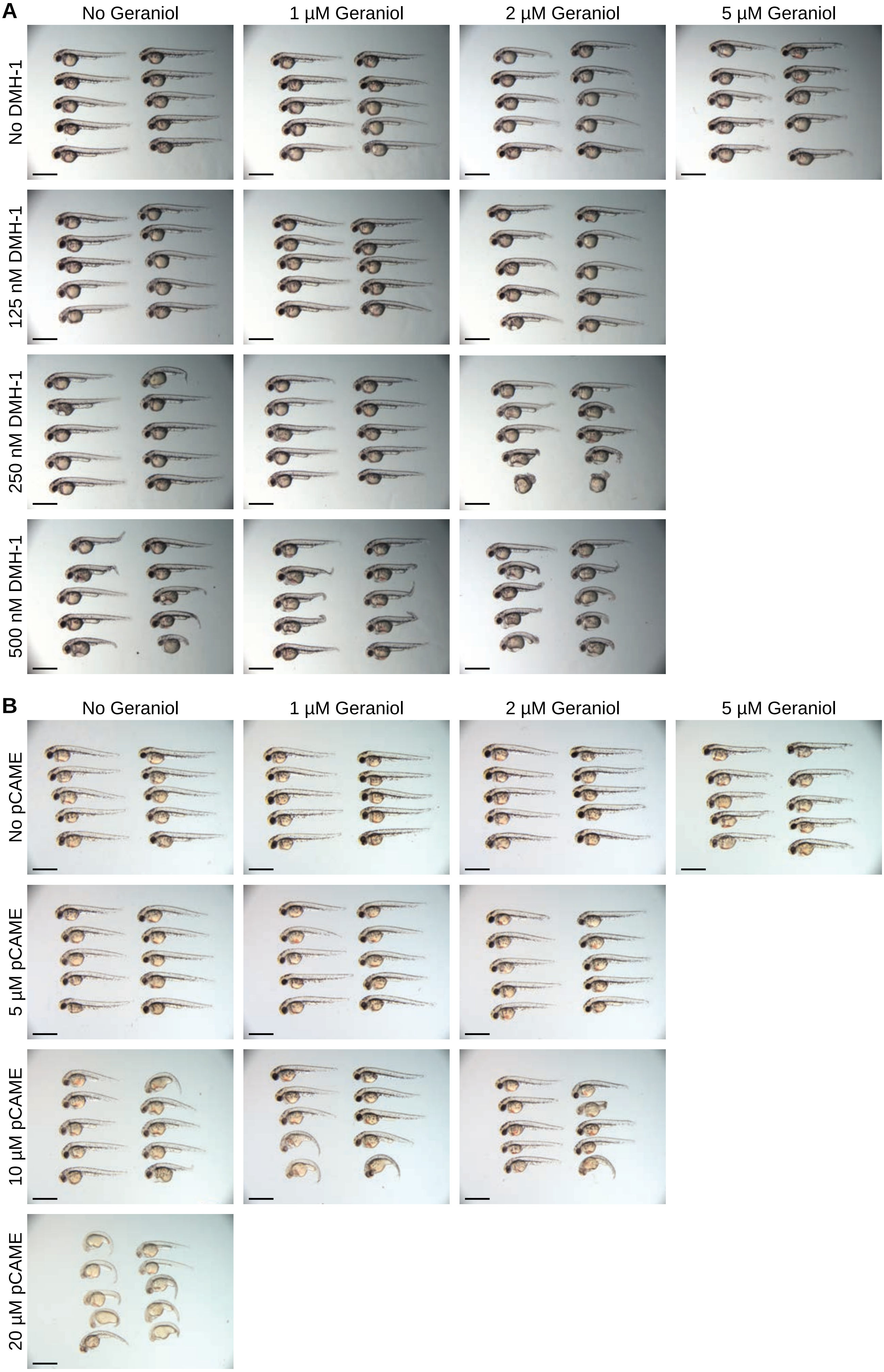
A
No Geraniol
1 µM Geraniol
2 µM Geraniol
5 µM Geraniol
No DMH-1
125 nM DMH-1
250 nM DMH-1
500 nM DMH-1
B
No Geraniol
1 µM Geraniol
2 µM Geraniol
5 µM Geraniol
No pCAME
5 µM pCAME
10 µM pCAME
20 µM pCAME

Fig S5

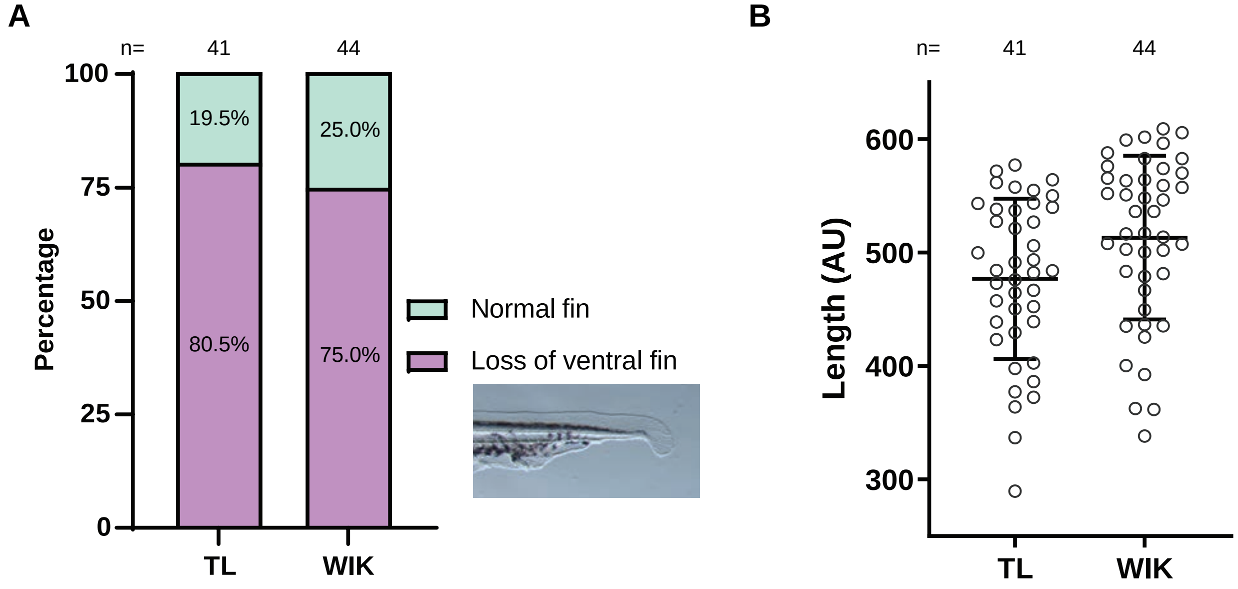
A
n=
41
44
100
75
50
25
0
Percentage
19.5%
25.0%
80.5%
75.0%
TL
WIK
Normal fin
Loss of ventral fin
B
n=
41
44
Length (AU)
600
500
400
300
TL
WIK

Fig S6

A

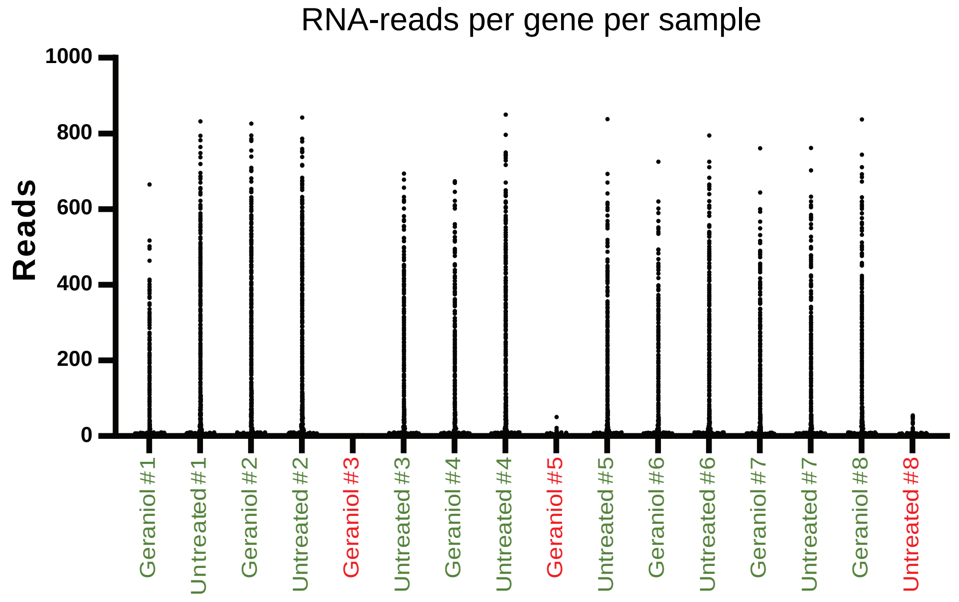

RNA-reads per gene per sample
Reads
1000
800
600
400
200
0
Geraniol #1
Untreated #1
Geraniol #2
Untreated #2
Geraniol #3
Untreated #3
Geraniol #4
Untreated #4
Geraniol #5
Untreated #5
Geraniol #6
Untreated #6
Geraniol #7
Untreated #7
Geraniol #8
Untreated #8


B

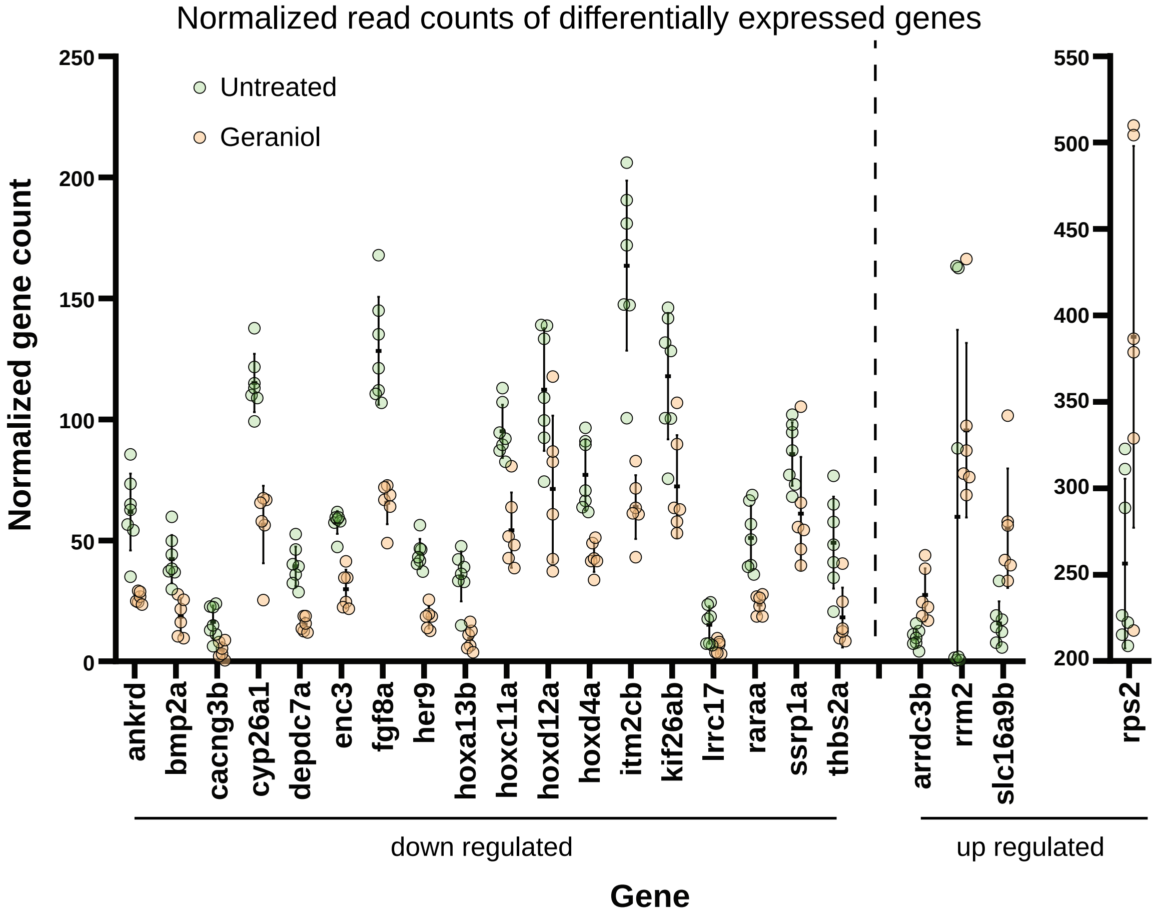

Normalized read counts of differentially expressed genes
Untreated
Geraniol
Normalized gene count
250
200
150
100
50
0
550
500
450
400
350
300
250
200
ankrd
bmp2a
cacng3b
cyp26a1
depdc7a
enc3
fgf8a
her9
hoxa13b
hoxc11a
hoxd12a
hoxd4a
itm2cb
kif26ab
lrrc17
raraa
ssrp1a
thbs2a
arrdc3b
rrm2
slc16a9b
rps2
down regulated
up regulated
Gene

Fig S7

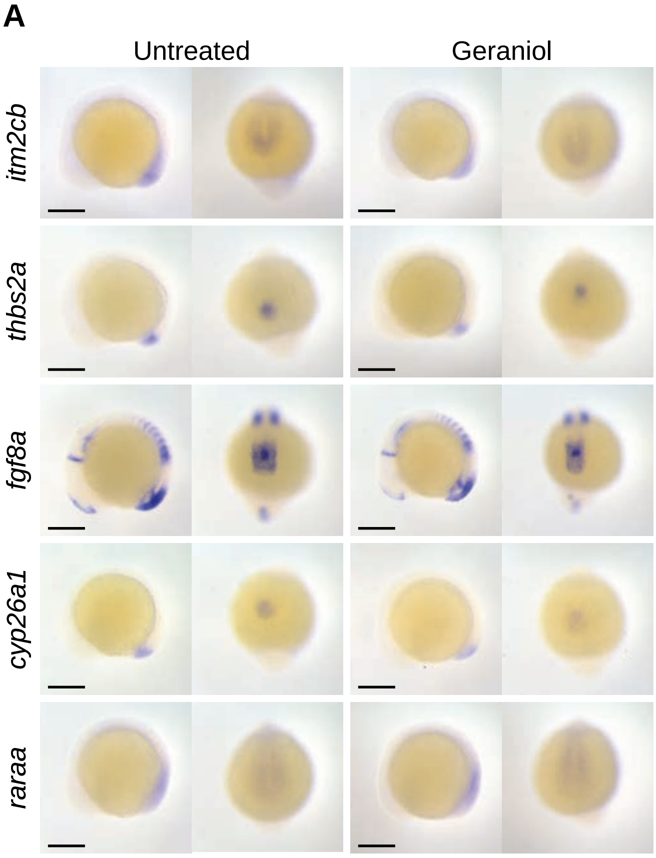
A
Untreated
Geraniol
itm2cb
thbs2a
fgf8a
cyp26a1
raraa

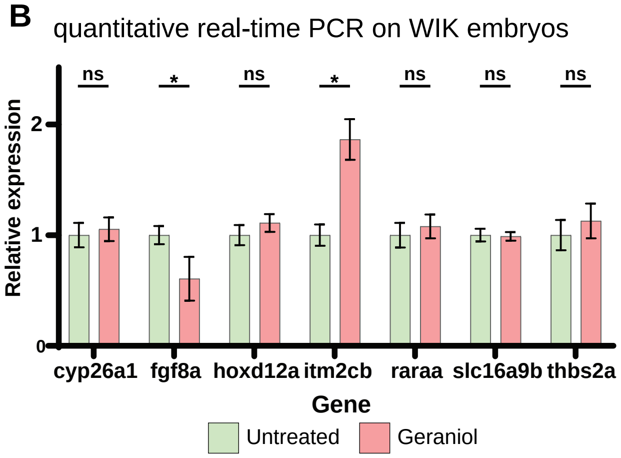
B
quantitative real-time PCR on WIK embryos
Relative expression
0
1
2
ns
*
ns
*
ns
ns
ns
cyp26a1
fgf8a
hoxd12a
itm2cb
raraa
slc16a9b
thbs2a
Gene
Untreated
Geraniol

Fig S8

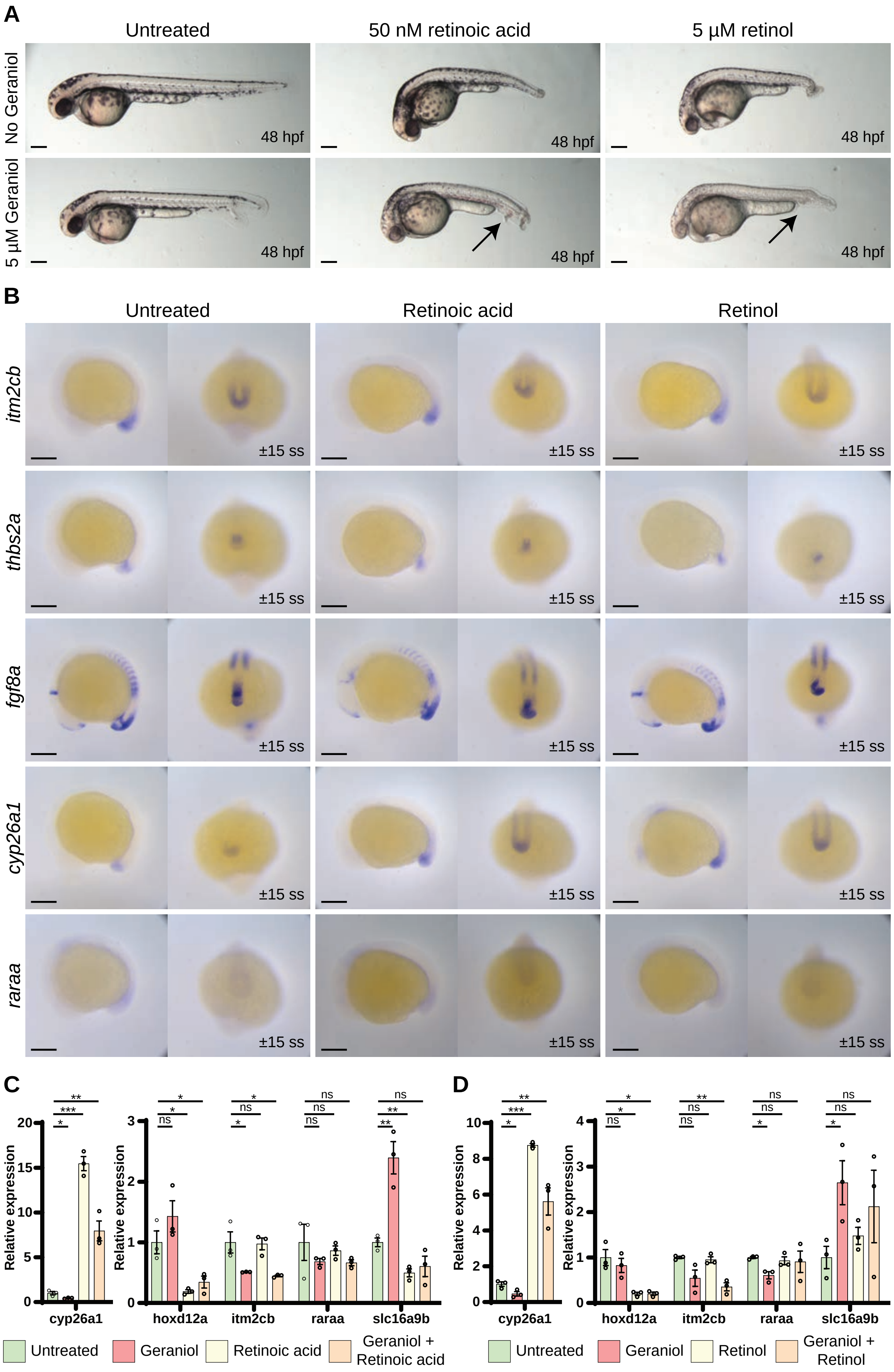
A
Untreated
50 nM retinoic acid
5 µM retinol
No Geraniol
5 µM Geraniol
48 hpf
B
Untreated
Retinoic acid
Retinol
itm2cb
thbs2a
fgf8a
cyp26a1
raraa
±15 ss
C
Relative expression
cyp26a1
hoxd12a
itm2cb
raraa
slc16a9b
Untreated
Geraniol
Retinoic acid
Geraniol + Retinoic acid
D
Relative expression
cyp26a1
hoxd12a
itm2cb
raraa
slc16a9b
Untreated
Geraniol
Retinol
Geraniol + Retinol

Fig S9

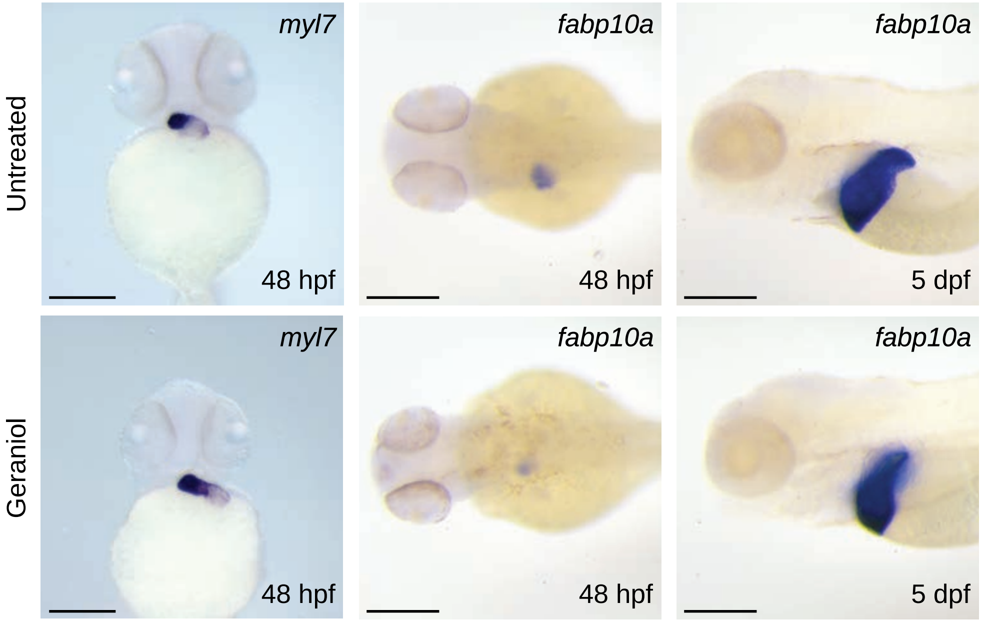
Untreated
Geraniol
myl7
fabp10a
fabp10a
48 hpf
48 hpf
5 dpf
myl7
fabp10a
fabp10a
48 hpf
48 hpf
5 dpf

**Table S1: primer sequences for probe synthesis**

| Gene | Forward primer sequence | Reverse primer sequence |
|---|---|---|
| *bmp2a* | AGCAGAGCCAACACTATCAG | GAG*TAATACGACTCACTATAGGG*TTAAGATGACCCGCTCGTAC |
| *col2a* | *from plasmid* | |
| *cyp26a1* | TCTACAAGACGCACCTCTTC | GAG*TAATACGACTCACTATAGGG*ACTTCTTTCATCGCCTGCAAA |
| *fabp10a* | GTGGCAGGTTTACGCTCAGG | GAG*TAATACGACTCACTATAGGG*GCTCTTCCTGATCATGGTGG |
| *fgf8a* | *from plasmid* | |
| *hoxd12a* | CCGCAGATGTCCTACAGTAG | GAG*TAATACGACTCACTATAGGG*AGTTCCAATCTGTCCGAAAGC |
| *itm2cb* | GTCTCCCTTCAGTGGCCTTT | GAG*TAATACGACTCACTATAGGG*GCGTCTCAGTCTGTAGGTCT |
| *msgn1* | CGTGAGGACAGGTCATTTGG | GAG*TAATACGACTCACTATAGGG*CGAGGATGCCGGATAACTCT |
| *myl7* | *from plasmid* | |
| *myod1* | TTCTACGACGACCCTTGCTT | GAG*TAATACGACTCACTATAGGG*TCCGTCTTCTCGTCTGACAC |
| *ngn1* | *from plasmid* | |
| *ntl* | *from plasmid* | |
| *raraa* | CCTGCCTCGACATACTGATT | GAG*TAATACGACTCACTATAGGG*AGAGCACACAGTAGTCCGTAA |
| *thbs2a* | CCGTCAGCTTGAGATAGTGT | GAG*TAATACGACTCACTATAGGG*CATGACCTGCCTCTTTGTTG |
| *xirp2a* | CAGTACCTCAAATGCTCAGG | GAG*TAATACGACTCACTATAGGG*AATATCTCTGCTTGCCTCCTC |

**Table S2: Quantitative PCR primer sequences**

| Gene | Forward primer sequence | Reverse primer sequence |
|---|---|---|
| *β-actin* | TTCTGGTCGGTACTACTGGTATTGTG | ATCTTCATCAGGTAGTCTGTCAGGT |
| *cyp26a1* | TGTTAATGATCCGACGAGTC | CACATTATCAGCTCCCATCA |
| *fgf8a* | TACAAACGCAGGACTTTACA | TCACCTTACTTTGCTCACTC |
| *hoxd12a* | CCAGAATATCAGACCAGCTT | ACCAGATTTTGACTTGCTGA |
| *itm2cb* | AAATCCAAATGCCGTACTCT | TATCTGTAGAGGCACACTGA |
| *raraa* | GAGTTCAGAAGAGATCGTCC | ATGATGCAGTTCTTCTCTCG |
| *slc16a9b* | TCATTGCTAGTCCCATCTGC | TGGCAAGTGATGGTTACAGT |
| *thbs2a* | TTGTGGCCAAGGGATCTATC | TGTACCGACAGACACAGATT |
